# A Branching Process Model of Evolutionary Rescue

**DOI:** 10.1101/2021.06.04.447075

**Authors:** Ricardo B. R. Azevedo, Peter Olofsson

**Affiliations:** Department of Biology and Biochemistry, University of Houston, Houston, Texas, U.S.A.; Department of Mathematics, Physics and Chemical Engineering, Jönköping University, Sweden

**Keywords:** evolutionary rescue, adaptation, extinction, beneficial mutations, branching process, population dynamics

## Abstract

Evolutionary rescue is the process whereby a declining population may start growing again, thus avoiding extinction, via an increase in the frequency of fitter genotypes. These genotypes may either already be present in the population in small numbers, or arise by mutation as the population declines. We present a simple two-type discrete-time branching process model and use it to obtain results such as the probability of rescue, the shape of the population growth curve of a rescued population, and the time until the first rescuing mutation occurs. Comparisons are made to existing results in the literature in cases where both the mutation rate and the selective advantage of the beneficial mutations are small.

## 1 Introduction

A declining population may be saved from otherwise inevitable extinction by the establishment of one or more beneficial mutations, a phenomenon known as evolutionary rescue [4, 6, 12, 17]. Such rescue may be a desirable outcome, such as recovery of an endangered species, or an undesirable one, such as cancer becoming resistant to chemotherapy. A typical feature of evolutionary rescue is a U-shaped population growth curve corresponding to initial decline, stabilization, and ultimate recovery. This behavior has been established in mathematical models [12, 17, 25] and observed in experiments on real populations [2, 5, 30].

Orr and Unckless wrote two seminal papers on the mathematics of evolutionary rescue [24, 25]. They considered a population that has experienced sudden environmental deterioration such that it can no longer sustain itself. They then analyzed the possibility that this endangered population is able to adapt sufficiently rapidly to avoid extinction. They took into account the contributions of both standing genetic variation and new mutations to adaptation. Using methods from the theory of branching processes in combination with Haldane’s [13] classic approximation of the probability of fixation, Orr and Unckless obtained, for the first time, several highly interesting formulas for, among other things, the probability that a population is rescued either by standing genetic variation or by a new mutation, the expected size of a rescued population at a given time, and the waiting time for the first rescuing mutation.

Crucially, Orr and Unckless assumed that selection is weak: the change in the environment reduces the mean absolute fitness of the population slightly below 1, and beneficial mutations raise it slightly above 1; or, in branching process parlance, they assumed that the endangered population is just barely subcritical, and the rescued population is just barely supercritical. They further assumed that mutation is weak relative to selection.

The results of Orr and Unckless [24, 25] have been extended in multiple ways through studies of different kinds of models (e.g., birth-death processes [22], time-inhomogeneous branching processes [35], Feller diffusion processes [23], population genetic models [1, 36]), making a variety of assumptions (e.g., density-dependent population regulation [22, 35], population structure [35], rescue by multiple mutations [23, 26], variable mutational effects [1, 23, 26], epistasis [1, 26], recombination [36], environmental deterioration [22]).

In the present work, we return to the branching process model analyzed by Orr and Unckless but make no assumptions about the strengths of selection or mutation. We establish general results that coincide with those of Orr and Unckless when selection and mutation are weak, but can differ significantly in other scenarios. We also derive new results on the number of independent beneficial mutations contributing to rescue. Strong selection is frequently observed in nature, specially as a result of human activity [28], although the extent to which strong selection contributes to evolutionary rescue is unclear. Furthermore, mutation rates are widely variable among species [21] and can be increased by both genetic and environmental stress [11, 33]. Thus, our results are of more than purely theoretical interest because endangered populations do not necessarily meet the assumptions of weak selection and weak mutation.

## 2 Model and Preliminaries

### 2.1 Branching Process

The population is modeled by a discrete-time branching process consisting of two types of individuals: wildtype individuals and individuals carrying a beneficial mutation. For simplicity, we refer to the latter as mutants. All individuals reproduce asexually, independently of each other, and experience hard selection [37]. Wildtype individuals can have offspring of both types, whereas mutants can only have offspring of their own type. All mutations are beneficial; there are no back mutations.

Denote by *X* the total number of offspring of a wildtype mother, where *X* is a nonnegative integer-valued random variable with mean *E*[*X*] = 1 – *r*, where *r* is the degree of maladaptation of a wildtype individual (see Table 1 for a complete list of variables and parameters). The absolute fitness of a wildtype individual is, therefore, 1 − *r*. We assume throughout that 0 < 1 − *r* < 1, that is, 0 < *r* < 1. Thus, a population composed entirely of wildtype individuals is subcritical and doomed to extinction in the absence of rescue^1^. The maladaptation is assumed to arise from an abrupt change in the environment [12, 17, 24, 25].

**Table 1:** Variables and parameters

| Symbol | Description |
| --- | --- |
| $X$ | Total number of offspring of a wildtype individual |
| $W$ | Number of wildtype offspring of a wildtype individual |
| $B$ | Number of mutant offspring of a wildtype individual |
| $\varphi(t)$ | Probability generating function (pgf) of $X$ |
| $F(t)$ | Joint pgf of the $W$ and $B$ offspring of a wildtype individual |
| $G(t)$ | Pgf of the $B$ offspring of a mutant individual |
| $r$ | Degree of maladaptation of a wildtype individual |
| $u$ | Beneficial mutation rate |
| $s$ | Beneficial effect of a mutation |
| $\mathcal{D}$ | Event of extinction of the population |
| $\mathcal{R}$ | Event of rescue of the population |
| $Z_n$ | Total number of individuals in the population at generation $n$ |
| $W_n$ | Number of wildtype individuals in the population at generation $n$ |
| $B_n$ | Number of mutant individuals in the population at generation $n$ |
| $q_w$ | Probability of extinction of a population starting from one wildtype individual |
| $q_b$ | Probability of extinction of a population starting from one beneficial individual |
| $p_w$ | Probability of rescue of a population starting from one wildtype individual |
| $p_b$ | Probability of rescue of a population starting from one beneficial individual |
| $P(\mathcal{R})$ | Probability of rescue |
| $P_{\text{new}}(\mathcal{R})$ | Probability of rescue from new mutations |
| $P_{\text{sv}}(\mathcal{R})$ | Probability of rescue from standing variation |
| $m_{ij}$ | Number of type-$j$ offspring of a mother of type $i$ |
| $M$ | Mean reproduction matrix with elements $m_{ij}$ |
| $K$ | Total number of beneficial mutations arising in a population |
| $K_S$ | Total number of beneficial mutations successful in rescuing a population |
| $T$ | Waiting time of the first appearance of a beneficial mutation |
| $T_S$ | Waiting time of the first appearance of a beneficial mutation successful in rescuing a population |

Each offspring of a wildtype individual may be mutant with probability *u* (the beneficial mutation rate) or wildtype with probability 1 – *u*, independently of other offspring. We denote the number of wildtype and mutant offspring by *W* and *B*, respectively, so that we have *X* = *W* + *B*. Conditioned on *X, W* and *B* have binomial distributions with success probabilities 1 – *u* and *u*, respectively:

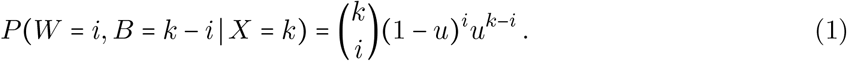

Mutant individuals may exist in the population as standing genetic variation or arise *de novo* through mutation. We assume a multiplicative fitness model throughout, that is, mutants have absolute fitness (1 – *r*)(1 + *s*), where *s* is the beneficial effect of the mutation they carry. Mutant individuals cannot accumulate additional beneficial mutations. We assume that *s* is large enough to make (1 – *r*)(1 + *s*) > 1. Thus, the process of mutant individuals is supercritical and has the potential to rescue the population from extinction.

Let *φ*(*t*) be the probability generating function (pgf) of *X*

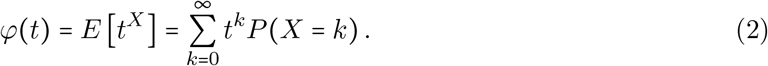

A complete specification of the pgf in Equation (2) requires that the distribution of *X* be known. To avoid uninteresting special cases, we make the natural assumption that *φ*(1) = *P*(*X* < ∞) = 1. Also, as *E*[*X*] < 1, we must have *φ*(0) = *P*(*X* = 0) > 0.

With *W* and *B* as above, let *P*(*i, j*) = *P*(*W* = *i, B* = *j*) and *P*(*j*) = *P*(*B* = *j*) be the offspring distributions for a wildtype and a mutant mother, respectively, and define the pgfs

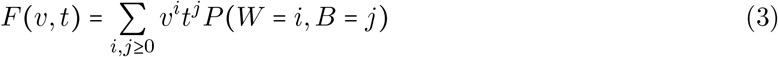

and

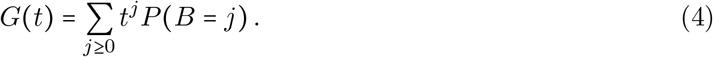

Obviously *φ*(*t*) = *F*(*t, t*) and for computations, the following result is of interest.

#### Lemma 2.1.

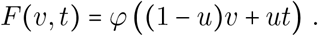

*Proof*.

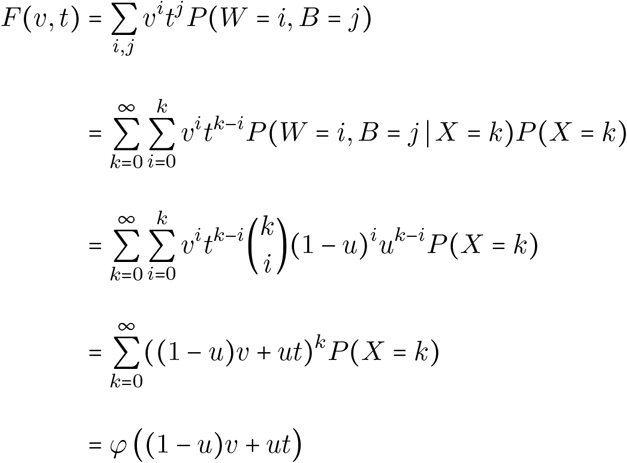

by the binomial theorem.

### 2.2 Extinction

Let *W_n_* and *B_n_* be the number of wildtype and mutant individuals, respectively, in generation *n*. The total population size is *Z_n_* = *W_n_* + *B_n_*. Let further 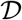 denote the event of extinction (death) of the population, that is

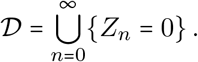

Denote by *q_w_* and *q_b_* the probabilities of extinction of the population when starting from one wildtype individual and one mutant individual, respectively:

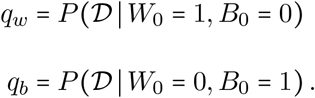

For convenience, we denote the corresponding rescue probabilities by

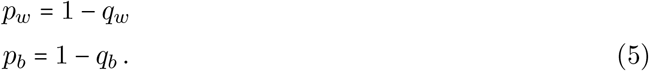

As the mutant type is supercritical we have *q_b_* < 1, and by standard branching process theory, *q_b_* can be found as the smallest solution in [0,1] to the equation *t* = *G*(*t*) (recalling that the process of mutant individuals is a single-type process)

Next we must find the extinction probability *q_w_* of a population that starts from one wildtype individual. As there is a chance of a beneficial mutation and mutants have a chance of avoiding extinction, we realize that *q_w_* < 1. It can further be computed explicitly according to the following result.

#### Proposition 2.2.

*Let ψ*(*t*) = *φ* ((1 – *u*)*t* + *uq_b_*). *The extinction probability q_w_ is the unique solution in* [0,1] *to the equation t* = *ψ*(*t*).

*Proof*. If *u* = 0, the equation reduces to *t* = *φ*(*t*), the usual equation for the extinction probability, and as 1 – *r* < 1, the unique solution is *t* = 1. Assume *u* > 0 and let 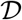 denote the event of extinction. Condition on the total number of offspring *X* and use the binomial theorem to obtain

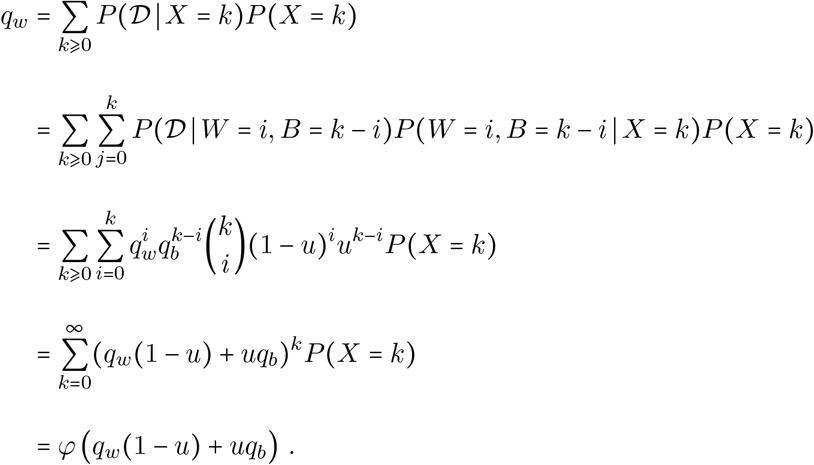

Thus *q_w_* solves the equation *t* = *ψ*(*t*). We next demonstrate that there is exactly one solution in [0,1] so the equation does indeed uniquely determine *q_w_*. As *φ* is strictly increasing and convex, so is *ψ*, and the claim follows if we can show that *ψ*(0) > 0 and *ψ*(1) < 1; the graph of *y* = *ψ*(*t*) must then intersect the line *y* = *t* exactly once in [0,1]. We note that

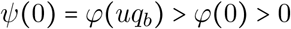

and

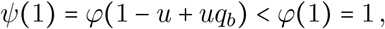

and the proof is complete.

### 2.3 Rescue

In our model a population may only experience one of two fates: extinction or rescue. Thus, if a population does not go extinct, we consider it rescued, and define the event 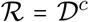, so that the probability of rescue is simply one minus the probability of extinction: 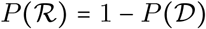. Hence, a population starting from *W*_0_ wildtype and *B*_0_ mutant individuals goes extinct if all the *W*_0_ + *B*_0_ independent subpopulations go extinct, and the probability of rescue is

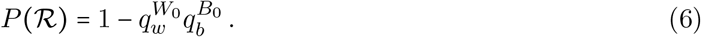

If initially the population is composed entirely of wildtype individuals (*B*_0_ = 0), rescue can only occur through new mutations. The probability that this kind of rescue occurs is given by (see Equation (6))

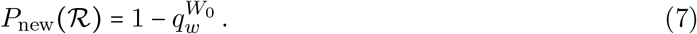

Alternatively, rescue can occur from the *B*_0_ mutant ancestors. This scenario is modeled by ignoring the contribution of new mutations, that is, *u* ≈ 0. In this case, *q_w_* = 1 because the wildtype population is subcritical, and Equation (6) becomes

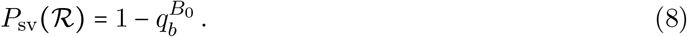

The results from this section enable us to compute the probability of rescue for any offspring distribution, any degree of wildtype maladaptation (*r*), any selective benefit of a mutation (*s*), any beneficial mutation rate (*u*), and any initial composition (*W*_0_, *B*_0_). Figure 1 illustrates the effect of some of these parameters on the probability of rescue from new mutations: the *P*_new_ increases with increasing *s* (Figure 1A), decreasing *r* (Figure 1B), and increasing *u* (Figure 1C and 1D). The results shown in Figure 1 were obtained assuming a Poisson offspring distribution. The observed effects can be expected to occur for other offspring distributions.

**Figure 1:**
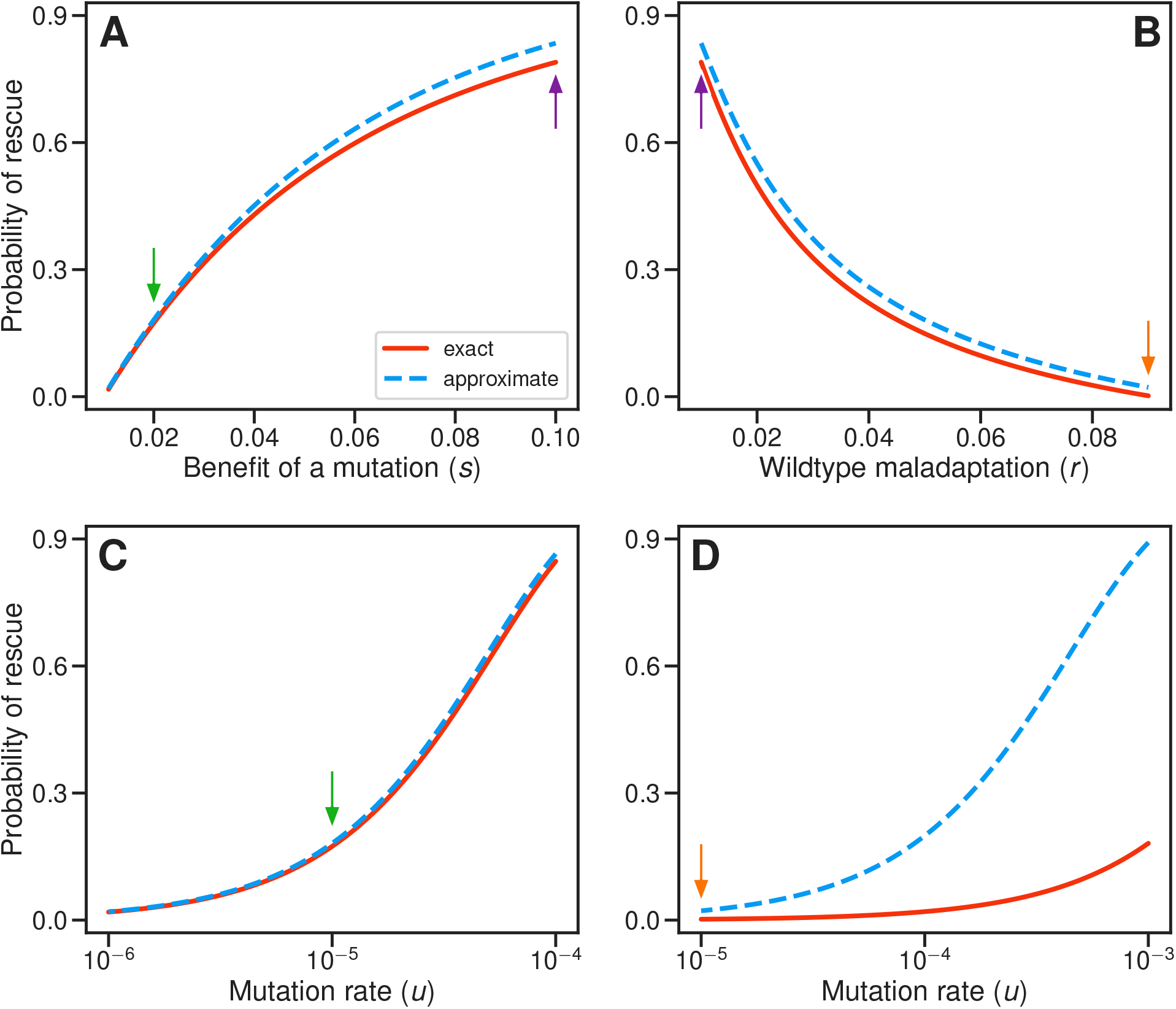
Effect of different parameters on the probability of rescue from new mutations, 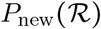. In all cases we assumed an initial population size of *W*_0_ = 10^4^ wildtype individuals (*B*_0_ = 0 mutant individuals). (A) Selection coefficient of a beneficial mutation, *s*. Other parameters: *r* = 0.01 and *u* = 10^−5^. (B) Degree of maladaptation of wildtype individuals, *r*. Other parameters: *s* = 0.1 and *u* = 10^−5^. (C–D) Beneficial mutation rate, *u*. Other parameters: (C) *r* = 0.01 and *s* = 0.02; (D) *r* = 0.09 and *s* = 0.1. Exact probabilities (red, continuous) were calculated using Equation (7), assuming a Poisson offspring distribution; *q_b_* and *q_w_* were calculated by solving the equations *t* = *G*(*t*) and *t* = *ψ*(*t*) numerically in [0,1], respectively. Approximate probabilities (blue, dashed) assuming weak selection/weak mutation were calculated using Equation (14). Arrows of particular colors indicate identical combinations of parameters.

In the next section we will consider the case of weak selection/weak mutation, that is, when *u, s*, and *r* are all small. The formulas given by Orr and Unckless [24, 25] arise as approximations in our general framework.

### 2.4 Weak Selection/Weak Mutation Approximation

Assuming weak selection, that is, sufficiently small *r* and *s*, we can get approximations of Equations (7) and (8) that express the survival probabilities in terms of *u, r, s, W*_0_, and *B*_0_. We also assume that *u* is small, by orders of magnitude, compared to *r* and *s* (i.e., weak mutation).

#### 2.4.1 Haldane approximation

Under weak selection, the expected number of offspring of a mutant individual is (1 – *r*)(1 + *s*) ≈ 1 + (*s* – *r*) and Haldane’s [13] approximation

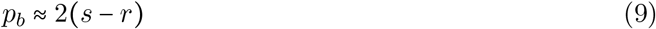

holds. We note here that although Haldane assumed a Poisson distribution, his approximation holds more generally by a first-order Taylor expansion of log *φ* (similar to what we do in Equation (10)). For distributions that, unlike the Poisson, have a variance unequal to its mean, there is an improved second-order approximation of *p_b_* [7]. For the purpose of this paper, however, we use the classical Haldane approximation.

#### 2.4.2 Standing variation

In the standing variation case with *u* ≈ 0, the probability of rescue in Equation (8) becomes

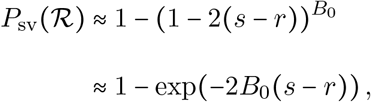

which is “*P_stand_*” in Equation (2) in Orr and Unckless [25].

#### 2.4.3 New mutations

For the new mutation case, a standard first-order Taylor expansion of the natural logarithm of *φ*(*t*) around *t* = 1 gives

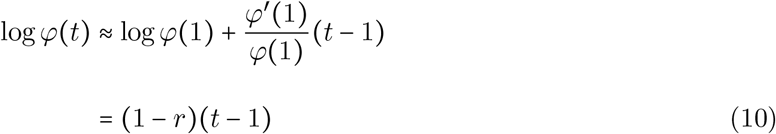

because *φ*′(1) = 1 – *r* and *φ*(1) = 1. (Note that for the Poisson offspring distribution, this Taylor approximation is exact.) Now recall

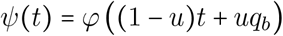

from Proposition 2.2. By Equation (10):

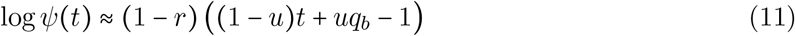

and the equation *t* = *ψ*(*t*) becomes

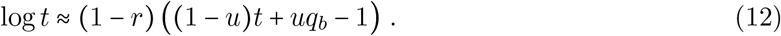

Using *q_b_* ≈ 1 – 2(*s – r*) and the first-order Taylor expansion log *t* ≈ *t* – 1, around *t* = 1, Equation (12) becomes

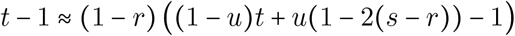

the solution of which is

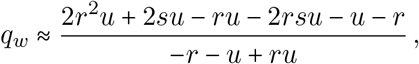

which gives

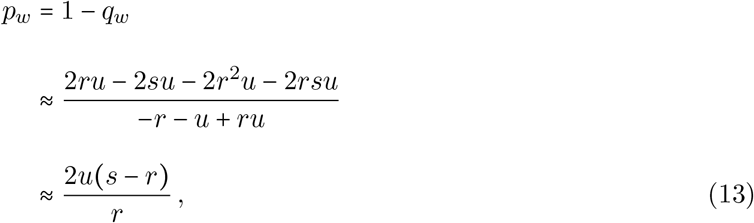

where the last approximation neglects the third-order terms 2*r*^2^*u* and 2*rsu* in the numerator, and *u* and *ru* in the denominator. Equation (7) becomes

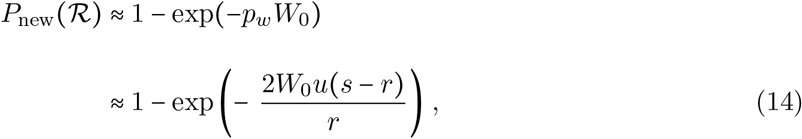

which is “*P_new_*” in Equation (3) in Orr and Unckless [24].

The weak selection/weak mutation approximation in Equation (14) performs well when the assumptions are met under a Poisson offspring distribution (Figure 1A, green arrow). However, it performs less well when selection is relatively strong (Figure 1A and 1B). In some scenarios, the approximation breaks down completely, even when the mutation rate is weak relative to selection (Figure 1D, high *u*). The reason for this discrepancy is that it relies on Haldane’s approximation (Equation (9)), which ignores a rs term. For example, using the parameters of Figure 1D (*rs* = 0.009) and *u* = 10^−3^ the probability of rescue according to Equation (14) is 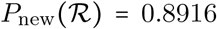, which is far from the exact value of 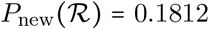 from Equation (7). But if we approximate the probability of fixation as *p_b_* = 2(*s – r – rs*), replace it in Equation (12), solve for *q_w_*, and use the full result without further approximations, we get 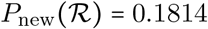.

## 3 Population Size

We next turn to the question of how the mean population size changes over time. Let the vector (*W_n_, B_n_*) be the number of wildtype and mutant individuals in generation *n*, and consider its expected value, the vector (*E*[*W_n_*], *E*[*B_n_*]). In standard branching process notation, let *m_ij_* denote the number of type-*j* offspring of a mother of type *i*. For simplicity, let the types be denoted by *w* and *b*, for wildtype and mutant, respectively. The mean reproduction matrix is

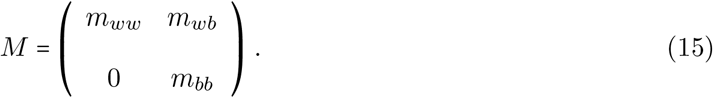

In our specific model, recall *E*[*X*] = 1 – *r*, the total number of offspring of a wildtype individual to get *m_ww_* = (1 – *r*)(1 – *u*), *m_wb_* = (1 – *r*)*u*, and *m_bb_* = (1 – *r*)(1 + *s*) so that

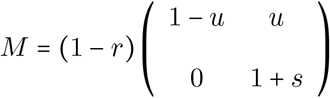

and it is readily seen by induction that its *n*th power is

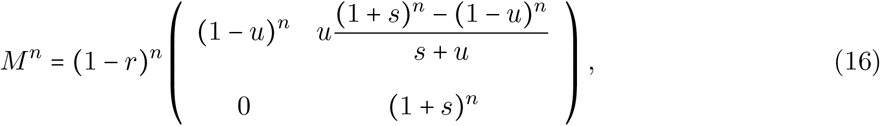

which grows ~ (1 – *r*)^*n*^(1 + *s*)^*n*^ as *n* → ∞, and the mutants ultimately dominate the population. Specifically, as *n* → ∞,

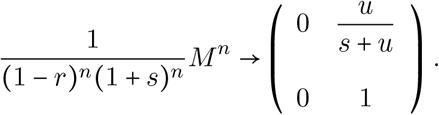

Starting from the vector (*W*_0_, *B*_0_) where *W*_0_ and *B*_0_ are fixed, we have

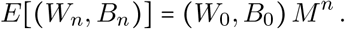

The total population size is *Z_n_* = *W_n_* + *B_n_*. Thus, the expected total population size, *E*[*Z_n_*] = *E*[*W_n_* + *B_n_*], starting from (*W*_0_, *B*_0_) ancestors is

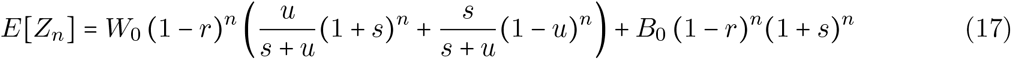

and asymptotically

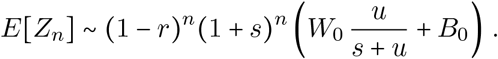

Equation (17) takes into account all populations, including those that go extinct. In section 5.1 we derive the expected population size of rescued populations.

## 4 Beneficial mutations

### 4.1 Number of Beneficial Mutations

Let 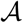 be the event that at least one beneficial mutation has arisen in a population started from one wildtype individual, and let 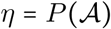. As above, let *φ* be the pgf of the number of offspring *X* (Equation (2)).

#### Proposition 4.1.

*The probability η is the unique solution to the equation*

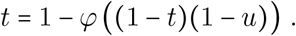

*Proof*. The right-hand side function is increasing in *t*. By putting in *t* = 0 and *t* = 1, respectively, we get the endpoint values 1 – *φ*(1 – *u*) and 1 – *φ*(0), both of which are in (0,1), and hence the right-hand side function intersects the line *y* = *t* exactly once in [0,1]. Next, condition on the number of offspring *X* of the ancestor, and the number *B* of those offspring that are mutants:

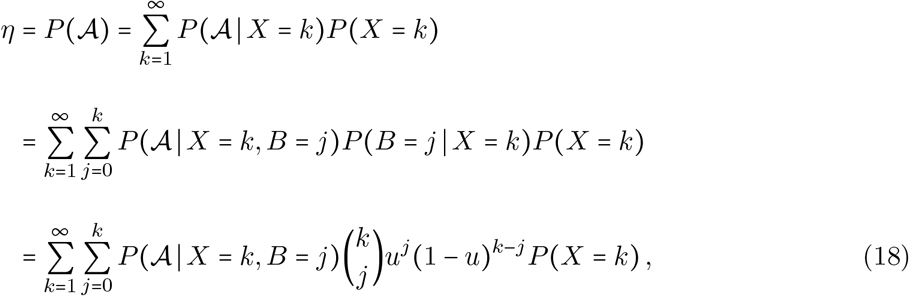

the sum starting at *k* = 1 because 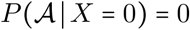. Next, note that we have 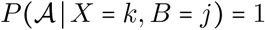 for *j* ⩽ 1, and for *j* = 0:

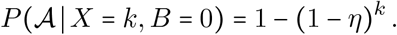

The term in Equation (18) with *j* = 0 equals

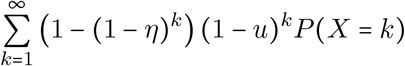

and the rest of the sum is

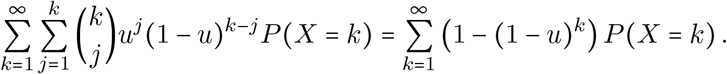

Add these together to get 1 – *φ*((1 – *η*)(1 – *u*)) and the proof is complete.

Note that, as there has to be at least one beneficial mutation for there to be rescue, we obviously have *η* > *p_w_*. Also note that with *υ* = 1 – *t*, the proposition may be rewritten as *υ* = *φ*(*υ*(1 – *u*)) where, by Lemma 2.1, the right-hand side equals the pgf of *W* integrated over the event that *B* = 0:

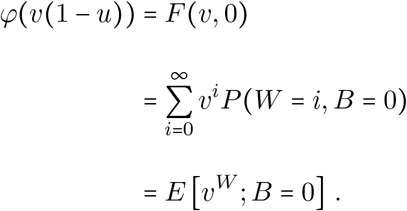

Next let *K* be the total number of beneficial mutations ever arising in a population. Its expected value is given by

#### Proposition 4.2.

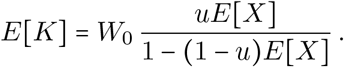

*Proof*. First let *W*_0_ = 1 and let *ν* = *E*[*K*]. We first rule out the possibility that *ν* = ∞. To this end, let *W* be the total number of individuals ever born in a branching process with mean *E*[*X*] = 1 – *r* < 1. Then

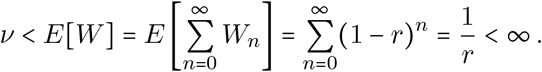

Next, condition on the number *X* of offspring of the ancestor, and the number *B* of those that are mutants. Noting that *E*[*K*|*X* = 0] = 0 we get

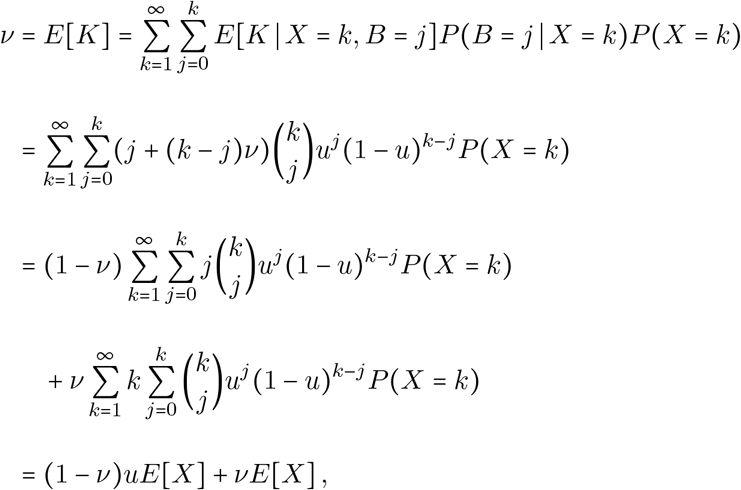

where we used that, for *j* ⩾ 0,

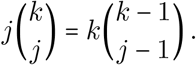

Solving *ν* = (1 – *ν*)*uE*[*X*] + *νE*[*X*] and multiplying by *W*_0_ concludes the proof.

Note that the proposition also follows from the informal argument that the expected number of mutant offspring of the ancestor is *uE*[*X*] and the populations starting from the remaining (1 – *u*)*E*[*X*] offspring produce on average *ν* new mutations each. Hence *ν* = *uE*[*X*] + (1 – *u*)*E*[*X*]*ν*, which also gives the result.

For example, with the parameters used in Figures 2A and 2B (*W*_0_ = 10^4^, *r* = 0.01, and *u* = 10^−5^), the expected number of independent beneficial mutations occurring during the course of evolution is *E*[*K*] = 9.9; the corresponding number for Figures 2C and 2D (*W*_0_ = 10^4^, *r* = 0.01, and *u* = 10^−4^) is *E*[*K*] = 98.0 (Proposition 4.2). These numbers of mutations include mutations that occur in populations that ultimately go extinct and mutations that occur in rescued populations but disappear and do not contribute to the rescue.

**Figure 2:**
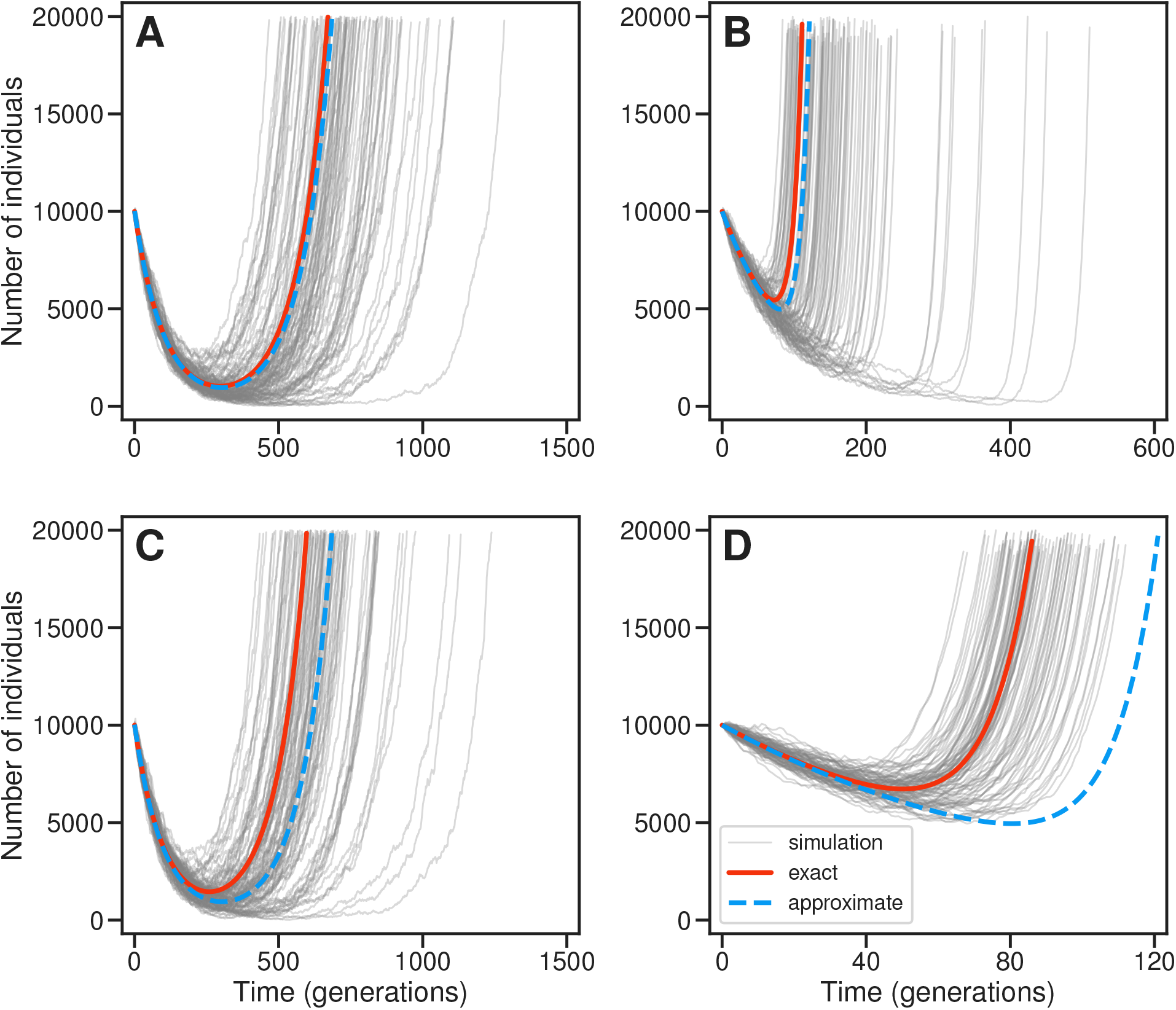
Total population size of rescued populations. We considered evolutionary rescue based on new mutations. In all cases we assumed an initial population size of *W*_0_ = 10^4^ wildtype individuals (*B*_0_ = 0 mutants), a degree of wildtype maladaptation of *r* = 0.01, and a Poisson offspring distribution. (A) Weak selection/weak mutation: *s* = 0.02 and *u* = 10^−5^ (green arrow, Figure 1A). (B) Stronger selection: *s* = 0.1 and *u* = 10^−5^ (purple arrow, Figure 1A). (C) Stronger mutation: *s* = 0.02 and *u* = 10^−4^ (far right, Figure 1C). (D) Stronger selection and stronger mutation: *s* = 0.1 and *u* = 10^−4^. Gray lines show total sizes (*Z_n_*) of individual simulated populations experiencing rescue (defined as reaching *Z_n_* > 2 × 10^4^). A sample of 100 individual trajectories is shown for each parameter combination. Exact population sizes (red, continuous) were calculated using Equation (24). Approximate population sizes (blue, dashed) assuming weak selection/weak mutation were calculated using Equation (26). The code for the simulations was written in Python 3.7 with NumPy version 1.21.0 [14] and is available at https://github.com/rbazev/rescue.

By Equation (5), a given mutation is successful in rescuing the population with probability *p_b_*. With *K_S_* denoting the number of successful mutations, we thus get

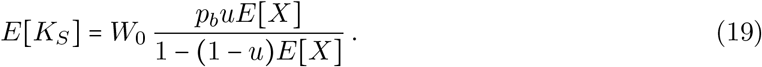

Equation (19) takes into account all populations, including those that go extinct (*K_S_* = 0). In section 5.3 we derive the *E*[*K_S_*] of rescued populations.

### 4.2 Waiting Time for a Beneficial Mutation

We next turn our attention to the time (generation) *T* of the first appearance of a beneficial mutation, aiming for the probabilities *P*(*T* > *n*), *n* = 0,1,2,…, in a population starting from wildtype individuals only. From these probabilities we can obtain the distribution function

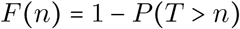

and the probability mass function

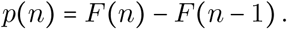

Note that *P*(*T* = ∞) > 0, because the population may die out without experiencing any mutations (specifically, *P*(*T* = ∞) = 1 – *η*, by Proposition 4.1). Thus, it also follows that *E*[*T*] = ∞.

We will establish an expression for *P*(*T* > *n*). To that end, first recall the pgf *φ*(*t*) of the number of offspring *X* of a wildtype mother (Equation (2)). Next, define a sequence of functions *H*_0_, *H*_1_,… recursively through

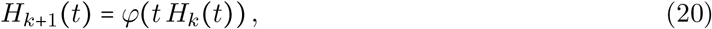

where *H*_0_(*t*) ≡ 1. Thus *H*_1_(*t*) = *φ*(*t*), *H*_2_(*t*) = *φ*(*tφ*(*t*)), and so on. Consider a population that starts from *W*_0_ wildtype individuals and no mutant individuals, *B*_0_ = 0.

#### Proposition 4.3.

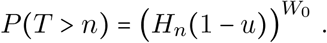

*Proof*. As *P*(*T* > *n*|*W*_0_ = *w*) = (*P*(*T* > *n*|*W*_0_ = 1))^*w*^, we can assume *W*_0_ = 1. Recall *W_n_* and *B_n_*, the number of wildtype and mutant individuals in generation *n*, respectively. Having *T* > *n* means that there are no mutation events at or before the *n*th generation, that is

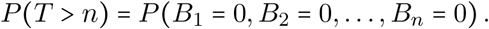

For any *i, j, k*, let

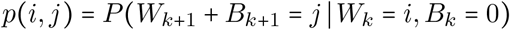

and note that

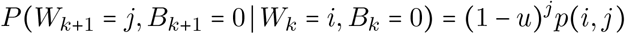

by Equation (1), and note also that

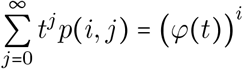

by elementary properties of pgfs. Repeated use of the above identities yields

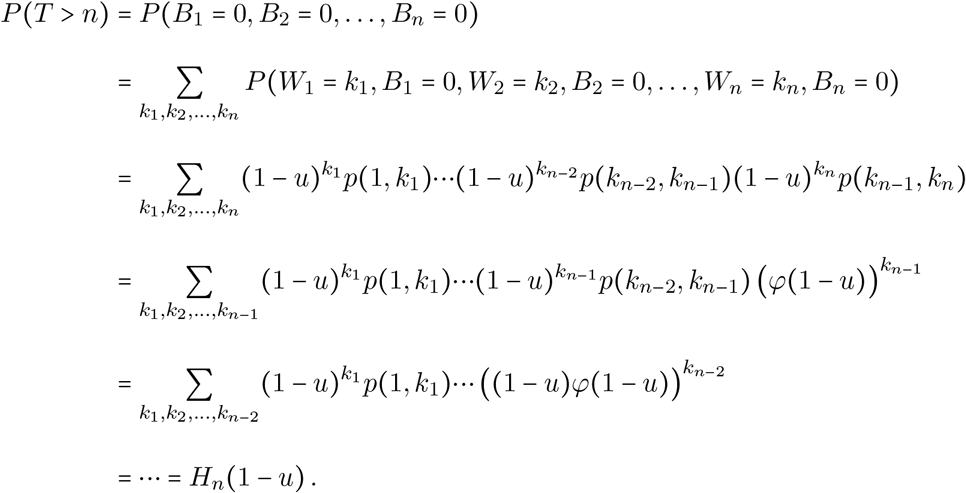

Recall that any given beneficial mutation is successful in rescuing the population with probability *p_b_*, and let *T_S_* be the time until such a successful mutation. The probabilities *P*(*T_S_* > *n*) can be found by applying Proposition 4.3 to the function 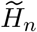, defined recursively through Equation (20) but instead using the pgf

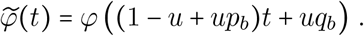

With *B_S_* and *B_F_* denoting the numbers of successful and failed mutants, respectively, 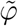 is the pgf of *W* + *B_S_* (recall Lemma 2.1). The total number of offspring is *X* = *W* + *B_S_* + *B_F_*, and conditioned on *X*, the three types of offspring follow a multinomial distribution with probabilities 1 – *u,up_b_*, and *uq_b_*. Thus, the conditional distribution of *B_S_* given *W* + *B_S_* is binomial with success probability

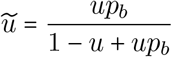

and we get

#### Corollary 4.3.1.

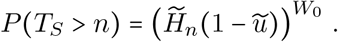

## 5 Rescued Populations

### 5.1 Population Size

For populations that survive, the mean population size (Equation (17)) is a poor measure as it takes into account all the populations that go extinct. For example, with an extinction probability of 99%, the 1% of populations that survive will have sizes far above the mean population size. Thus, we consider instead the mean population size conditioned on rescue. The analysis hinges upon use of the elementary result that for any integrable random variable *Y*, we have

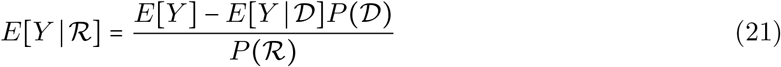

by the Law of Total Expectation.

By standard branching process results [3, 19], a supercritical branching process conditioned on extinction is a subcritical branching process, whose reproduction law can be given explicitly. To that end, recall the notation *P*(*i, j*) = *P*(*W* = *i, B* = *j*) for a wildtype mother and *P*(*j*) = *P*(*B* = *j*) for a mutant mother, and the corresponding pgfs *F*(*υ, t*) and *G*(*t*) (Equations (3) and (4)). Recall also the extinction probabilities *q_w_* and *q_b_*, starting from one wildtype and one mutant individual, respectively. Following Jagers and Lageras [19], the process conditioned on extinction has reproduction law given by

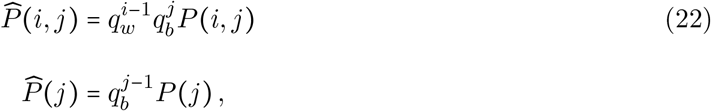

with the corresponding pgfs

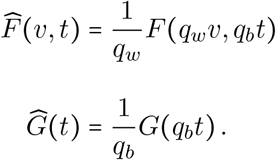

By standard properties of pgfs, we can compute the entries of the mean reproduction matrix 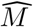, in analogy with Equation (15), as

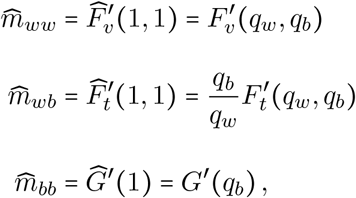

in the usual notation for partial derivatives. Hence

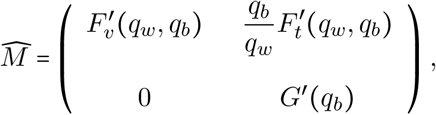

the *n*th power of which is

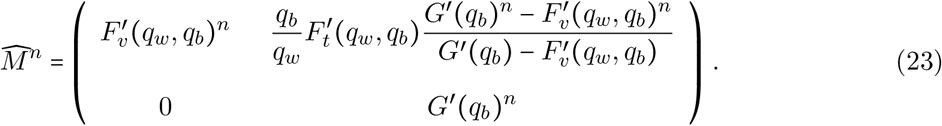

From Equations (21) and (23), we can now compute the *n*th generation mean population sizes conditioned on rescue, starting from *W*_0_ wildtype and *B*_0_ mutants as

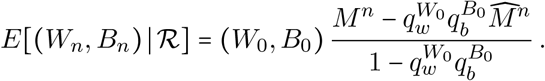

The total population size is *Z_n_* = *W_n_* + *B_n_* and we get

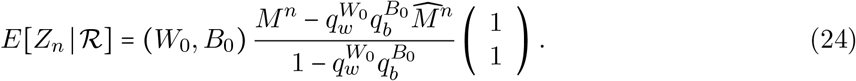

### 5.2 Weak Selection/Weak Mutation Approximation

If we assume that *u, r*, and *s* are all small we can get approximations of Equation (24) for the cases where rescue occurs through either standing variation or new mutations.

#### 5.2.1 Standing variation

We first consider the standing variation case where rescue is due to pre-existing mutant individuals and not to new mutations (*u* ≈ 0). Let the population start from (*W*_0_, *B*_0_) individuals where *B*_0_ is small. We then have

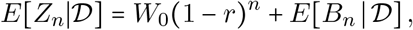

where, by Equation (4)

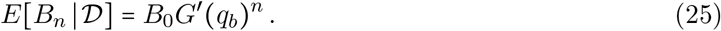

As

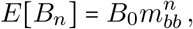

Equations (16), (21) and (25) yield

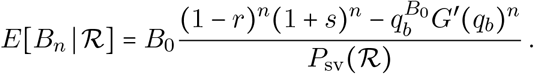

If *q_b_* is close to 1 and *B*_0_ small, we get the approximation

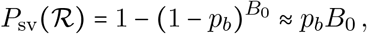

which yields

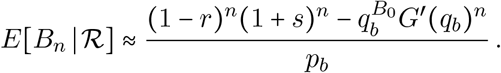

With Haldane’s approximation (Equation (9)) we get

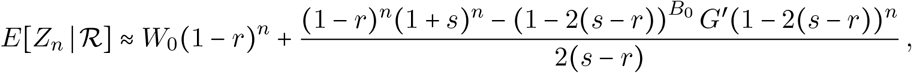

which is essentially Equation (10) in Orr and Unckless [25], noting that they use the approximation (1 – *r*)^*n*^(1 + *s*)^*n*^ ≈ *e*^(*s–r*)*n*^, and neglect the term 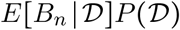 in Equation (21).

#### 5.2.2 New mutations

For the new mutation case, the population is rescued by mutation and not by pre-existing mutants. We thus let *B*_0_ = 0 and start from *W*_0_ wildtype individuals. Again, neglect the term 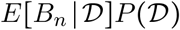 in Equation (21) to obtain

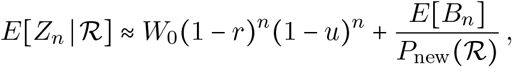

and Equation (16) gives, for the second term,

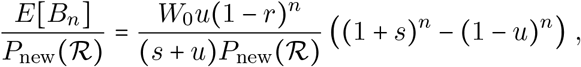

where

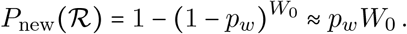

Neglecting the term (1 – *u*)^*n*^ and using Equation (13) to approximate *p_w_*, we now get

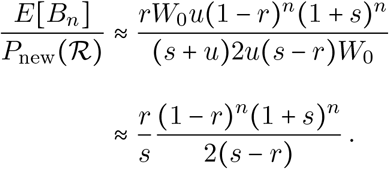

And, therefore,

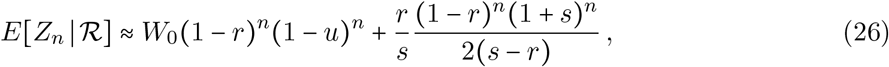

in agreement with Equation (19) in Orr and Unckless [25].

Figure 2 compares Equations (24) and (26) to individual-based stochastic simulations of the branching process under different strengths of selection and mutation. The weak selection/weak mutation approximation can perform reasonably well when selection is strong (Figure 2B), but can break down even when the mutation rate is weak relative to selection (Figure 2D). The weak selection/weak mutation approximation tends to underestimate population size (Figure 2) while overestimating the probability of rescue (Figure 1). This is because Equation (26) depends on 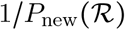, which leads to an underestimate of 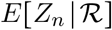 if 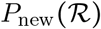 is overestimated.

### 5.3 Number of Rescuing Mutations

In Equation (19), we established an expression for the expected number of successful mutations, *E*[*K_S_*], in a population. By noting that

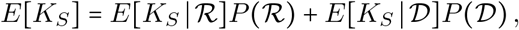

where, obviously, 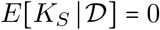, we get the following expression for the expected number of rescuing mutations in a rescued population starting from *W*_0_ wildtype individuals:

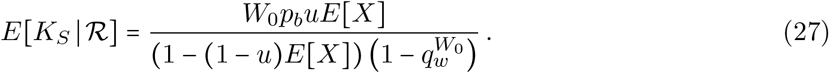

As there must be at least one rescuing beneficial mutation in a rescued population, we realize that 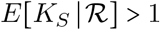.

For example, with the parameters used in Figure 2A (*W*_0_ = 10^4^, *r* = 0.01, *s* = 0.02, and *u* = 10^−5^), the expected number of independent rescuing mutations occurring during the course of evolution is 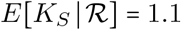 (Equation (27)). The corresponding numbers for the other parts of Figure 2 are: 2.0 (B), 2.2 (C), and 15.6 (D).

### 5.4 Waiting Time for a Rescuing Mutation

We next turn our attention to the times *T* and *T_S_* of the first appearances of a beneficial mutation and a successful mutation, respectively, in a rescued population. Proposition 4.3 and Corollary 4.3.1 give the unconditional probabilities *P*(*T* > *n*) and *P*(*T_S_* > *n*), and the Law of Total Probability gives

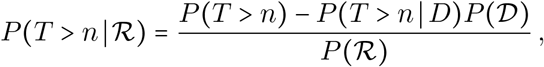

with a similar expression for *T_S_*. Note that conditioning on rescue implies that there is at least one mutation event in finite time, that is, we have 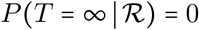. In contrast, both 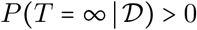 and *P*(*T* = ∞) > 0, because the population may die out without experiencing any mutations. Therefore, *E*[*T*] = ∞ and 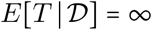, whereas 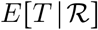 may still be finite (which also means that we cannot apply Equation (21) to find 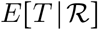).

Proposition 4.3 shows how to compute the probability *P*(*T* > *n*). To get to 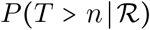 we need to compute 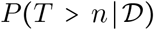 by use of the conditional offspring law 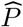 of Equation (22), but alas, there is no immediate analog of Proposition 4.3. This proposition relies upon the conditional distribution of mutants being binomial when conditioned on the total number of offspring; this property does not transfer to 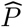 (which can be realized by simple examples, for example binary splitting). However, the case of a Poisson offspring law does lend itself to such analysis, which we address in the next section.

Things are easier when it comes to *T_S_*, which is also the more interesting variable, answering the question when the (first) rescuing mutation arises. As extinction obviously means that there are no successful mutations, we have 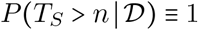 and hence

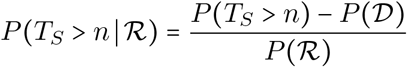

and by Corollary 4.3.1 we get

#### Corollary 5.0.1.

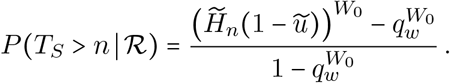

We can now obtain the probability mass function

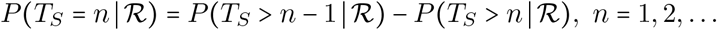

and the expected value

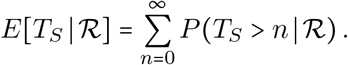

Orr and Unckless [25] provided the following weak selection/weak mutation approximation (their Supporting Information Text S2):

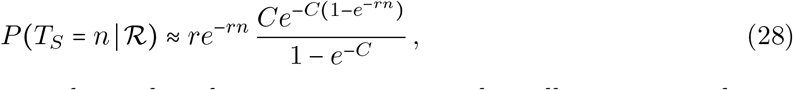

where *C* = 2*W*_0_*u*(*s – r*)/*r*. Figure 3 shows that this approximation works well, even in conditions where the approximation for population size breaks down (compare Figures 2D and 3D).

**Figure 3:**
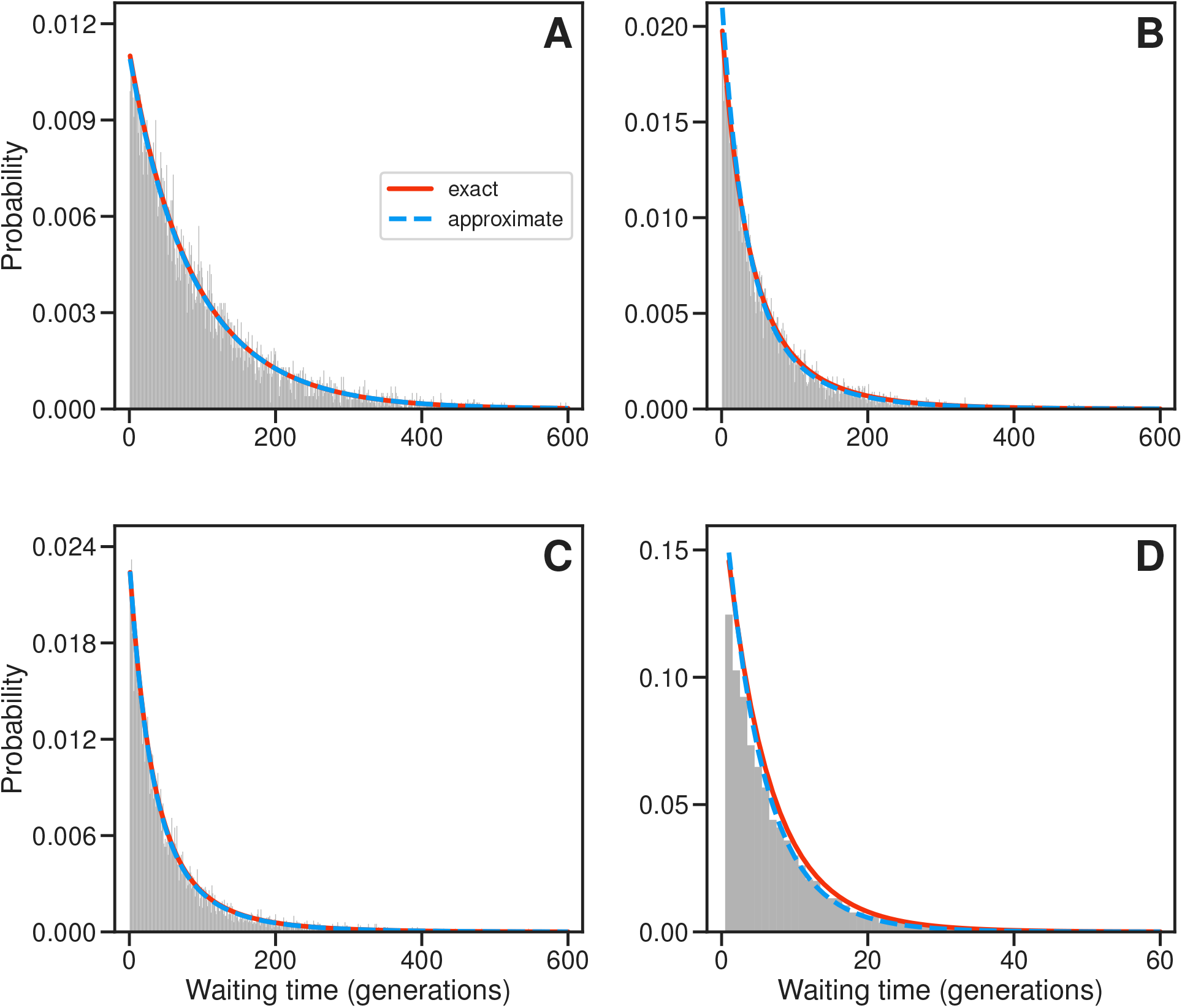
Waiting time for the first rescuing mutation. We considered evolutionary rescue based on new mutations. The parameter combinations in each part as the same as in Figure 2. In all cases we assumed an initial population size of *W*_0_ = 10^4^ wildtype individuals (*B*_0_ = 0 mutants), a degree of wildtype maladaptation of *r* = 0.01, and a Poisson offspring distribution. (A) Weak selection/weak mutation: *s* = 0.02 and *u* = 10^−5^ (green arrow, Figure 1A). (B) Stronger selection: *s* = 0.1 and *u* = 10^−5^ (purple arrow, Figure 1A). (C) Stronger mutation: *s* = 0.02 and *u* = 10^−4^ (far right, Figure 1C). (D) Stronger selection and stronger mutation: *s* = 0.1 and *u* = 10^−4^. Histograms of waiting times are based on 10^4^ stochastic simulations. The histograms are truncated for clarity, which leads to the exclusion of < 3% of values in each case. Rescue was defined as reaching *Z_n_* > 1.5 × 10^4^. Exact probabilities (red, continuous) were calculated using Corollary 5.0.1. Approximate probabilities (blue, dashed) assuming weak mutation were calculated using Equation (28).

### 5.5 The Poisson Distribution

As the Poisson distribution is frequently used as offspring distribution (as in our simulations), we finish with a section about its properties in the context of evolutionary rescue. By standard properties of the Poisson distribution, the numbers of wildtype and mutant offspring, *W* and *B*, are independent with *W* ~ Poi((1 – *r*)(1 – *u*)) and *B* ~ Poi((1 – *r*)*u*). As it turns out, both the Poisson distribution and the independence still hold when conditioning on the population going extinct. By Equation (22) and independence, the reproduction law conditioned on extinction is

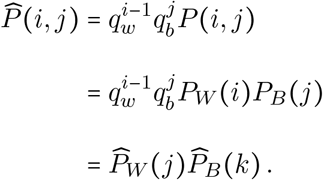

The last equality follows from the multiplicative separation of *i* and *j* in the expression for 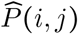; by this, the marginal distributions are independent. Hence 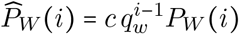 where the normalizing constant *c* is obtained by summing over *i*:

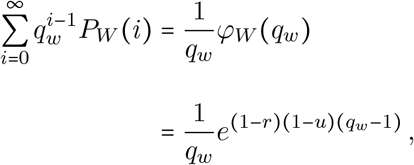

yielding

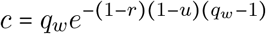

and thus

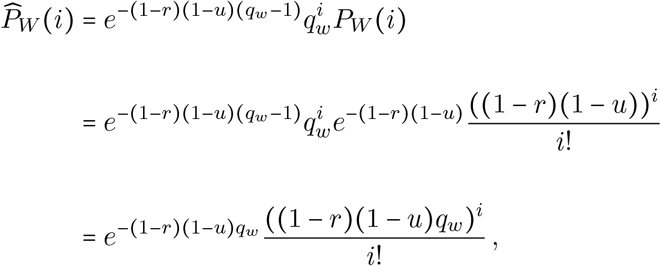

which we recognize as a Poisson distribution with mean (1 – *r*)(1 – *u*)*q_w_* and may write as

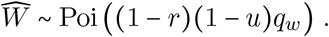

A similar calculation yields

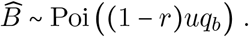

Put together, these also give the total number of offspring

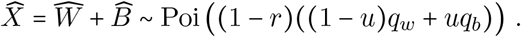

A mutant has a number of (necessarily mutant) offspring *B* which is Poi((1–*r*)(1+*s*)). Conditioned on extinction, more similar calculations give

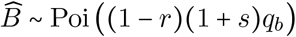

for a mutant mother. By standard properties of the Poisson distribution, we also get the conditional distributions of 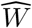 and 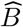 given 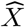 as binomial 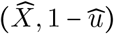 and 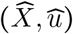, respectively, where

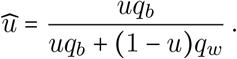

As *q_w_* > *q_b_*, we have 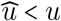 which makes sense.

The standard Poisson properties do not, however, carry over to the population conditioned on rescue. By the Law of Total Probability:

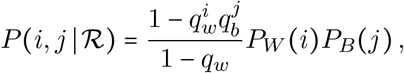

showing that, conditioned on rescue, *W* and *B* are neither Poisson nor independent.

## 6 Discussion

We use a two-type branching process to model evolutionary rescue introduced by Orr and Unckless [24, 25]. The types are wildtype individuals and individuals carrying beneficial mutations. The parameters are: the degree of maladaptation of wildtype individuals (*r*), the selective benefit of a mutation (*s*), the beneficial mutation rate (*u*), and the initial size and composition of the population (*W*_0_, *B*_0_). Our model is simple, but provides a basis for an understanding of the interplay between fundamental parameters in evolutionary rescue. Our formulas for probability of rescue, the expected size of a rescued population, and the waiting time for the first rescuing mutation boil down to those of Orr and Unckless [24, 25] in cases of weak selection/weak mutation, but the discrepancy can be large when selection is strong (Figures 1D and 2D). We derive new expressions for the number of independent beneficial mutations contributing to the rescue of a population.

Strong selection is frequently observed in nature, notably when humans are involved [28]. Examples include global climate change [10] and the use of antibiotics [27] and pesticides [15]. Strong levels of maladaptation (i.e., high *r*) are common. For example, shifting a population of the cowpea seed beetle, *Callosobruchus maculatus*, adapted to mung bean to lentil, caused approximately 99% mortality [31]. This extreme maladaptation caused multiple experimental populations to go extinct, but one population adapted quickly and reached 69% survival in 5 generations. High mutation rates are also common [21]. Furthermore, mutation rates have been shown to increase under precisely the kinds of stressful conditions that may put populations at risk and, therefore, in need of rescue [11, 33]. Thus, our results are of more than purely theoretical interest.

Application of our model to real populations is not straightforward because the mutational parameters (*s* and *u*) are difficult to estimate. The baseline values we adopted in our numerical exploration of the model (*s* = 0.02 and *u* = 10^−5^; e.g., green arrow in Figure 1A) are consistent with estimates obtained in three independent experimental studies on populations of the bacterium *Escherichia coli* [16, 29, 34]. Stronger mutational effects and a lower mutation rate (*s* ≈ 0.1 and *u* ≈ 10^−7^) were observed in a different study on the same species [38]; weaker mutational effects and a higher beneficial mutation rate (*s* ≈ 0.01 and *u* ≈ 10^−4^) were observed in the yeast *Saccharomyces cerevisiae* [8]. Thus, the scenarios in the numerical illustrations in Figures 1–3 involve plausible values of the mutational parameters.

Our model makes the simplifying assumption that all beneficial mutations have the same effect. Studies in both *S. cerevisiae* [8] and the bacterium *Pseudomonas fluorescens* [18] found evidence that the beneficial effects of mutations are approximately exponentially distributed. A study in two different bacteriophages (ID11 and *ϕ*6) found support, instead, for a uniform distribution of mutational effects [32]. Our model could be extended to consider more than one type of mutant individual [1, 23].

Another central assumption of our model is that an individual can acquire at most one beneficial mutation. This assumption precludes rescue involving multiple mutations in our model. There is experimental evidence that evolutionary rescue through individuals carrying multiple mutations does occur in real populations. For example, populations of *S. cerevisiae* “rescued” from high copper concentrations often acquired multiple beneficial mutations [9]. Osmond et al. [26] showed that if beneficial mutations of small effect are more common than those of large effect then rescue by multiple mutations can be more likely than rescue by single mutations.

Most models of evolutionary rescue [12, 24, 25], including ours, ignore deleterious mutations. This modeling decision is questionable because deleterious mutations are expected to be orders of magnitude more likely to occur than beneficial ones. The accumulation of deleterious mutations is predicted to accelerate the decline in population size in a process known as mutational meltdown [20]. Thus, the probability of rescue from new mutations predicted in a model like ours is likely to be overly optimistic.

Our model makes several additional simplifying assumptions including, but not restricted to, a constant environment, asexual reproduction, and density-independence. The influence of these assumptions on the probability, tempo, and mode of evolutionary rescue is an active area of research. Different studies have incorporated, for example, a deteriorating environment [22], recombination [36], and density-dependent population regulation [22, 35].

## Acknowledgements

We thank Alex Stewart and Daniel Weissman for helpful discussions and Logan Chipkin for his early contribution to this project. The National Science Foundation grants DEB-1354952 and DEB-2014566 (awarded to R.B.R.A.) and National Institutes of Health grant R15GM093957 (awarded to P.O.) funded this work.

## Footnotes

1 Many of the results in the paper are valid also in the critical case 1 − *r* = 1, that is, *r* = 0.

## Notes

https://github.com/rbazev/rescue

### Competing Interest Statement

The authors have declared no competing interest.

### Summary of Updates

Revision in response to a round of peer-review.

